# Contextual Regression: An Accurate and Conveniently Interpretable Nonlinear Model for Mining Discovery from Scientific Data

**DOI:** 10.1101/210997

**Authors:** Chengyu Liu, Wei Wang

**Affiliations:** Department of Chemistry and Biochemistry, University of California San Diego, La Jolla, CA 92093-0359; Department of Cellular and Molecular Medicine, University of California San Diego, La Jolla, CA 92093-0359

## Abstract

Machine learning algorithms such as linear regression, SVM and neural network have played an increasingly important role in the process of scientific discovery. However, none of them is both interpretable and accurate on nonlinear datasets. Here we present contextual regression, a method that joins these two desirable properties together using a hybrid architecture of neural network embedding and dot product layer. We demonstrate its high prediction accuracy and sensitivity through the task of predictive feature selection on a simulated dataset and the application of predicting open chromatin sites in the human genome. On the simulated data, our method achieved high fidelity recovery of feature contributions under random noise levels up to ±200%. On the open chromatin dataset, the application of our method not only outperformed the state of the art method in terms of accuracy, but also unveiled two previously unfound open chromatin related histone marks. Our method fills in the gap of accurate and interpretable nonlinear modeling in scientific data mining tasks.

## Introduction

Predictive models are important tools for data analysis. Linear models, such as linear^1^ and logistic regression^2^, have been essential prediction methods for a very long time. They can provide adequate prediction accuracy while their linear formulation makes them suitable for inferring relationship between prediction target and features. Recently, complex nonlinear models have gained increasing popularity as an alternative to linear models. Methods such as kernel SVM^3^, neural network^4^ and decision trees^5^ can achieve better prediction performance than the linear models especially on nonlinear datasets. In biology, for instance, deep convolutional neural network (DCNN) methods such as DeepBind^6^ and DeepSea^7^ have been developed for scanning the genome for regions of interest with state of the art accuracy.

This accuracy improvement, however, also comes with a cost: the parameters of a nonlinear model are hardly human interpretable. This not only makes model improving and debugging difficult but also prevents us from acquiring knowledge from these models and validating their findings, which is particularly important when controversial conclusions can be reached from the prediction results^8,9^. While examining the filters^10,11^ in a DCNN can offer some insight of the important feature combinations, it cannot provide quantification of the feature contributions: in the later layers, the features are processed through multiple rounds of matrix multiplication, addition and neuron activation which makes their contributions to the output intractable. Besides, this approach cannot be applied to other neural networks or machine learning methods such as feedforward neural network (FNN)^12^ or Long Short Term Memory (LSTM)^13^ neural network. To fill this gap, many well-thought and intriguing methods such as DeepLift^14^, LIME^15^ and SHAP^16^ have been developed for quantifying feature contributions in general machine learning models. All of them are based on the same overall strategy. When using these methods, the user first needs to choose a reference data point with the target value of interest, a genomic region with enhancer label of value 1 for example. Then the program will generate permuted data points by making small changes to the input features, which forms an interrogating dataset. The interrogating dataset are fed into a trained model of user's interest and the output of the model are collected. From the magnitude of the output change caused by the input change, an interpreting model that describes the feature contribution to the prediction result can be generated for that data point. This approach, however, has a few practical problems. First, since interpretation is based on each user selected reference data point, it is inherently a “local” model and thus can be overfitted to that point. Second, the training and interpretation processes are decoupled, which blocks the possibility of real time model monitoring during training. Third, the user is required to be familiar with the dataset in order to pick representative reference points, which is not an easy task particularly in scientific data when (i) the prediction target value is a continuous value other than binary; (ii) the dataset contains natural noise that causes the target value to deviate from its real value; (iii) the data are of large quantity and diversity.

We have developed a new method called contextual regression to address these challenges. It can quantify feature contributions during training without user intervention while preserving the accuracy achieved by a complex nonlinear model: we train the model to learn an embedding function to map each feature vector to a corresponding linear model that can predict the target value most accurately (Figure 1). In principle, values of the elements in the input feature vector describe its context and the embedding serves as a classifier of the context. It generates a continuous value vector as the class of the context which is similar to the mechanism of attention^17^ and word2vec^18^. We call this class of the context “context weight” and thus this method the “contextual regression.” In this way, the contribution of each feature can be inferred from the statistics of the context weights on the dataset. We demonstrated its accuracy and high interpretability through quantifying feature contributions to prediction accuracy in a simulated dataset with known “ground truth” and an application to the open chromatin prediction.

**Figure 1.**
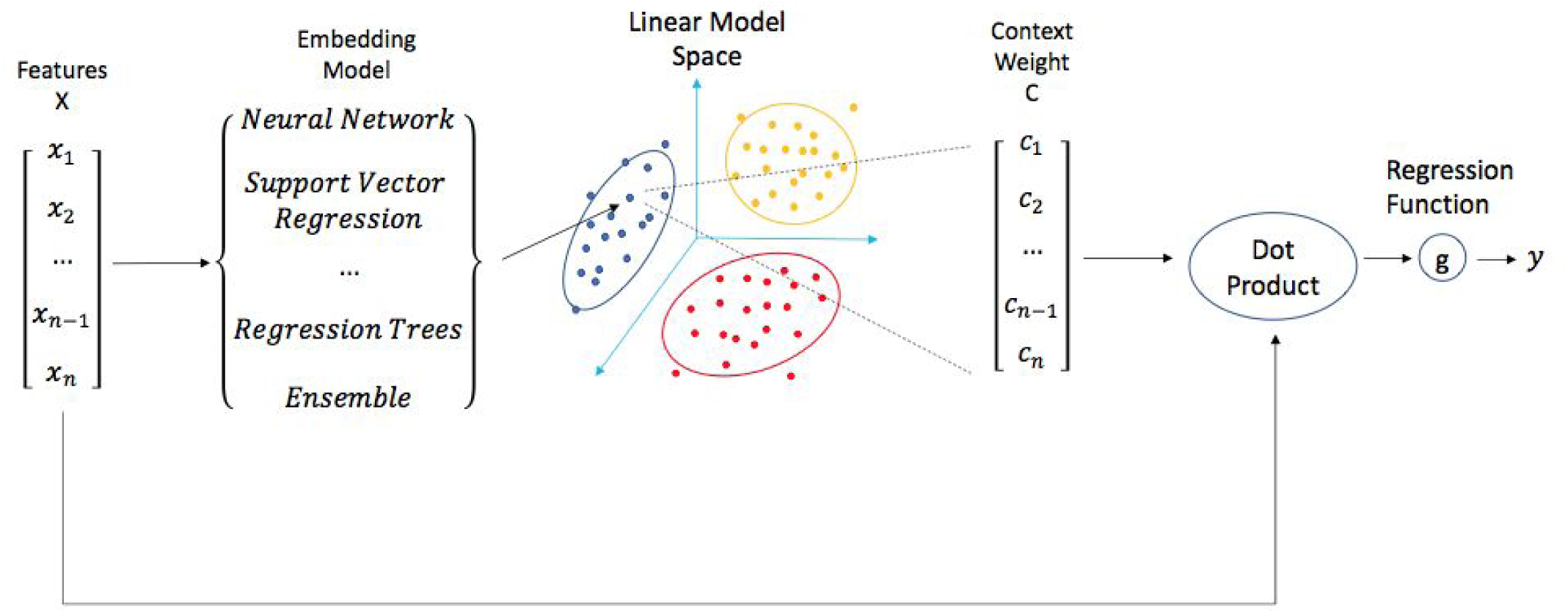
A graphic demonstration of the contextual regression model. Dot product apply the linear model, which is the context weight vector C, to the feature vector X. The embedding model can be any machine learning methods that can produce a vector output from a vector input. Different colors of points in the linear model space is an illustration that they can be classified into different subtypes.

## Results

### The contextual regression method

As shown in Figure 1, our model aims to learn a pseudo-linear model that approximates the function:

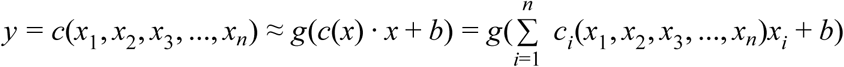

where y is the prediction target, x_i_ are the features, b is a constant, g is the regression function (for example, it is an identity function for linear regression and is a logistic function for logistic regression), and c is the embedding function, which calculates the context weight from the feature vector x as the linear model for the prediction of y at that data point.

Thus, rather than trying to fit all the data with a single linear model, our method learns many linear models and apply different ones at different data points according to their values. Like illustrated in Figure 1, the embedding network takes a data point (represented as a vector) as its input, and outputs a linear model (the context weight, represented as a vector). Then the data point is dot-producted with the linear model to output the prediction. This method, in its property, can handle nonlinear relationship to make accurate predictions while still producing human-readable linear relationship between features and the prediction target. One potential caveat, an old problem in predictive modeling, is the existence of alternative model on the dataset, i.e., very different model parameters can yield similar prediction accuracy. Since a descent direction for y is not necessarily a descent direction for each individual *c*_*i*_ (see supplement for detail), this scenario can possibly happen. We have developed the utopia penalty technique to address it. This technique reveals equivalent or interchangeable features by adding a term to the cost function that penalizes unbalanced context weights. The setup and an application example of it can be found in the supplement.

We evaluated the performance of the proposed method using a simulated dataset and a DNaseI hypersensitivity (DHS) dataset^19^. In both cases, the features have distal relationship and thus we used the bidirectional-Long Short Term Memory (LSTM) neural network, a specialized neural network model for distally related data, to be the main component of our embedding function.

### Evaluating the Contextual Regression model on simulated data

The simulated dataset includes an artificial “ground truth” in its feature-target relationship that we wish to find. We sampled features x_i_ from an exponential distribution to (1) reduce the possibility of alternative models such that we can test the rule extraction ability of our method, and (2) imitate the scenario that the feature values differ in order of magnitude such as the sequencing read count in biological data. In the simulated data, we consider 50 features x_i_ that are in sequential order. Each element *c*_*i*_ of the context weight vector C is determined by the value of x_i_ and its neighbors. The “ground truth” context weights *c*_*i*_ and then the target value y are calculated from x_i_ using the formula below. The context weight formula is chosen such that it is composed of common functions observed in natural data (linear, square and square root) and includes relationship with the neighboring elements. More details about the setup can be found in the supplement.

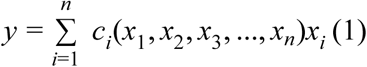

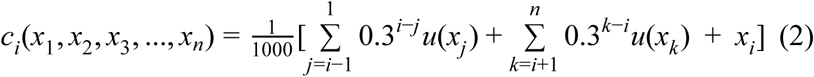

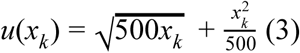

Our model was tested under 5 noise levels from ±0% to ±80%. When adding the noise, a random number inside the noise range was sampled uniformly (for instance, between −80% to +80% for noise level ±80%) and this fraction was added or subtracted from the target value y. The model was trained on 70% of the randomly selected data (training set) and then its fitness was tested on the remaining 30% data (testing set). We used root mean square error (RMSE) as the measure of prediction accuracy. Cosine distance measures the angle difference between two vectors, and thus is a good measure of vector similarity. So for the measure of rule extraction fidelity, we used root mean square cosine distance (RMSCD), the cosine distance version of RMSE, between the “ground truth” weights and the weights produced by our embedding network at their corresponding data points. Since the target values contain noise, it is impossible for the model to achieve 100% accuracy. So we compared RMSE with the expected error under the corresponding noise level, which was calculated by integrating the percent error in the error range. The details of this calculation can be found in the supplement.

Under all noise levels, our model achieved high performance in both prediction accuracy and rule extraction fidelity on the testing set (Figure 2 and 1S), with RMSE less than 1% away from the expected error and RMSCD below 0.03. (Table 1S). We also visually inspected (Figure 2) some randomly selected “ground truth” context weights (green) vs context weights (blue) produced by the embedding network under noise level ±0% and ±80% (Figure 2). They are indeed visually similar which agrees with the overall low RMSCD values between the “ground truth” and network output context weights.

**Figure 2.**
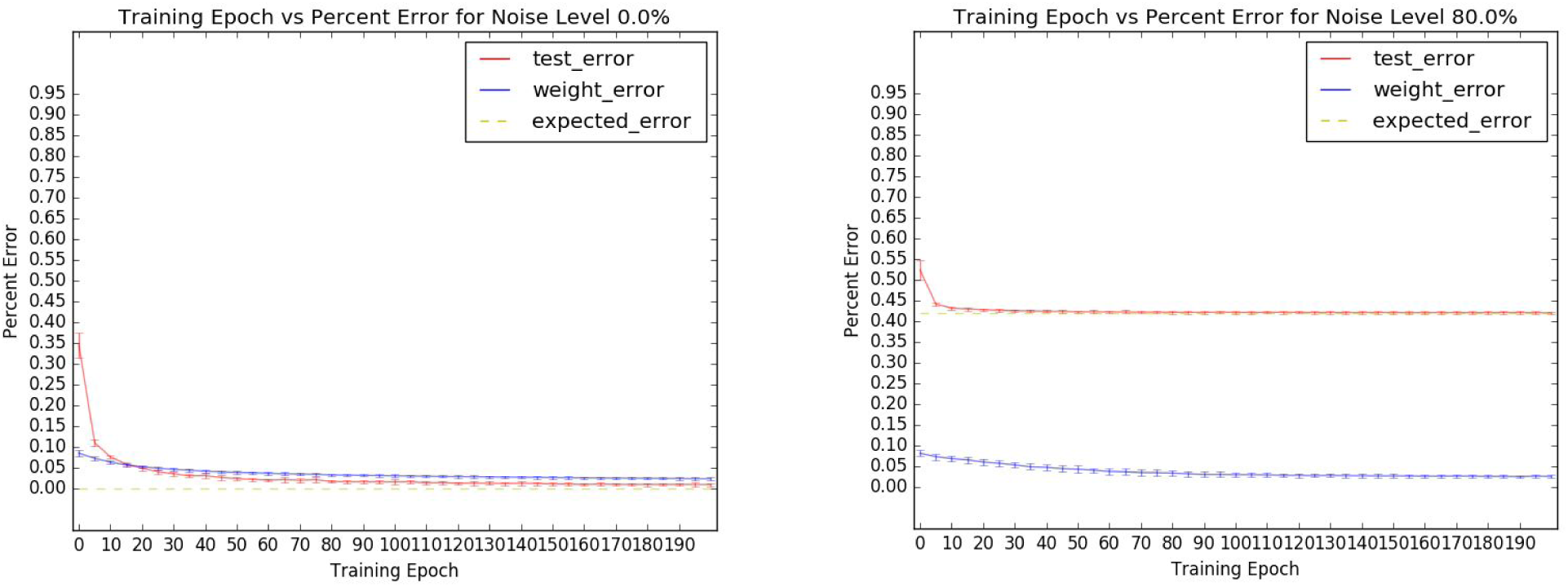
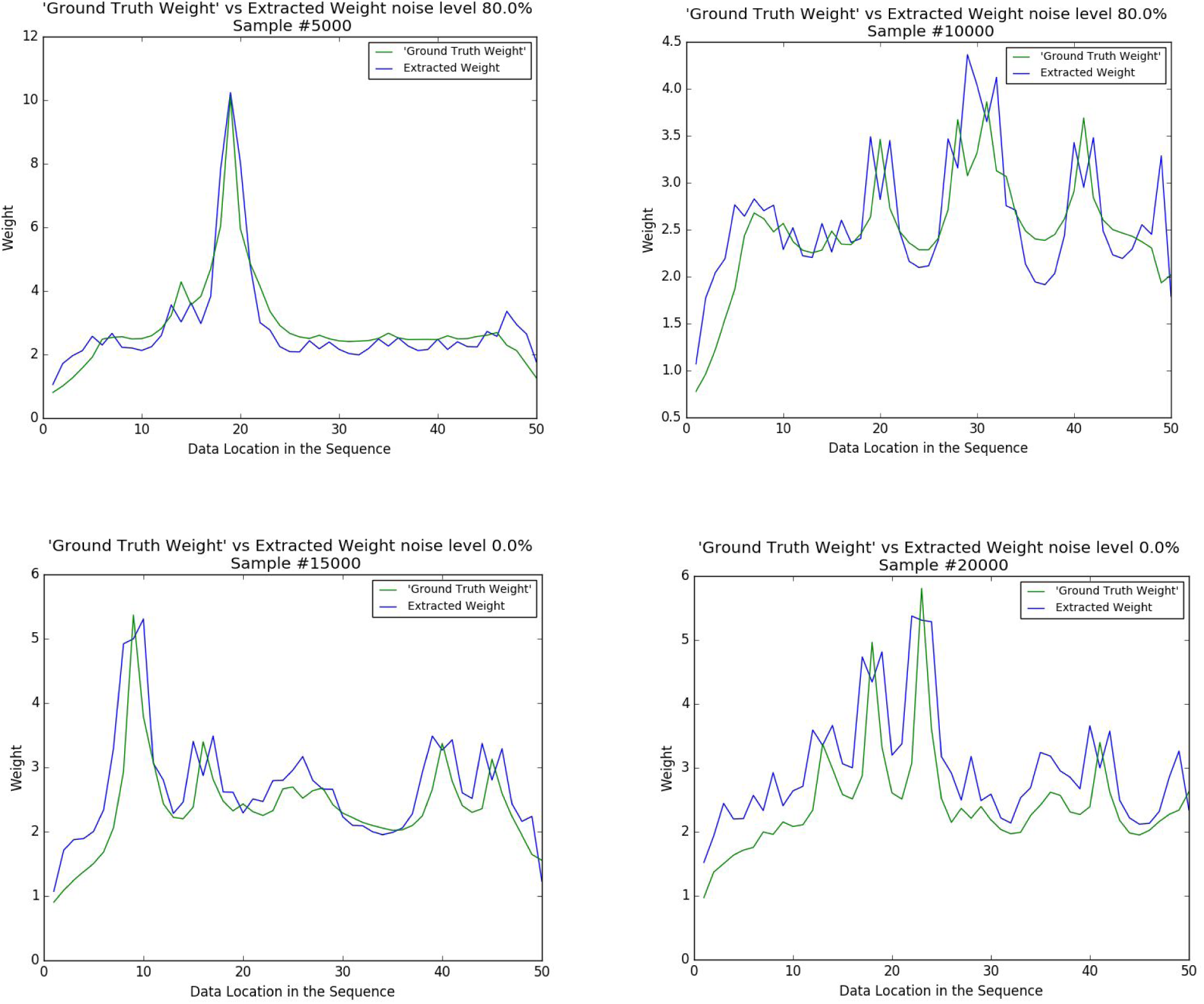
Performance assessment of contextual regression on simulated dataset- Top: Contextual regression performance on the testing set under noise level 0% and 80%. Height of the error bar is 1 standard deviation of RMSE or RMSCD among 20 runs. In each epoch (training cycle), the model was trained on 10% of the randomly selected data from the training set and tested on the whole testing set. We chose 10% of the data as one epoch, other than 100% that is ordinarily used, to show the evolution of error. Middle: Sample plot of “ground truth” context weight (green) vs context weight (blue) calculated by the embedding net in the contextual regression model for ±80% noise dataset. Bottom: Same comparison plots for ±0% noise dataset.

In certain scientific experiments, the signal to noise ratio can be extremely low. This is especially the case in some experiments of physics^20,21^, social science^22,23^ and biological science^24–26^. Thus we also tested our model under noise level of 200% (signal to noise ratio 1:2), 500% (signal to noise ratio 1:5) and 1000% (signal to noise ratio 1:10). We found that under all three noise levels, the prediction errors still approach the expected error very fast (Figure 2S and Table 2S). However, the RMSCD is only stably decreasing under noise level of 200% and 500%. Under 1000% noise, the RMSCD value starts to slightly increase and fluctuate when the training time gets longer. This is confirmed in our visual inspection of the sample data points (Figure 3S), where the sampled extracted weights still resembles the corresponding “ground truth” weights under 200% and 500% noise but distorted under 1000% noise.

**Figure 3.**
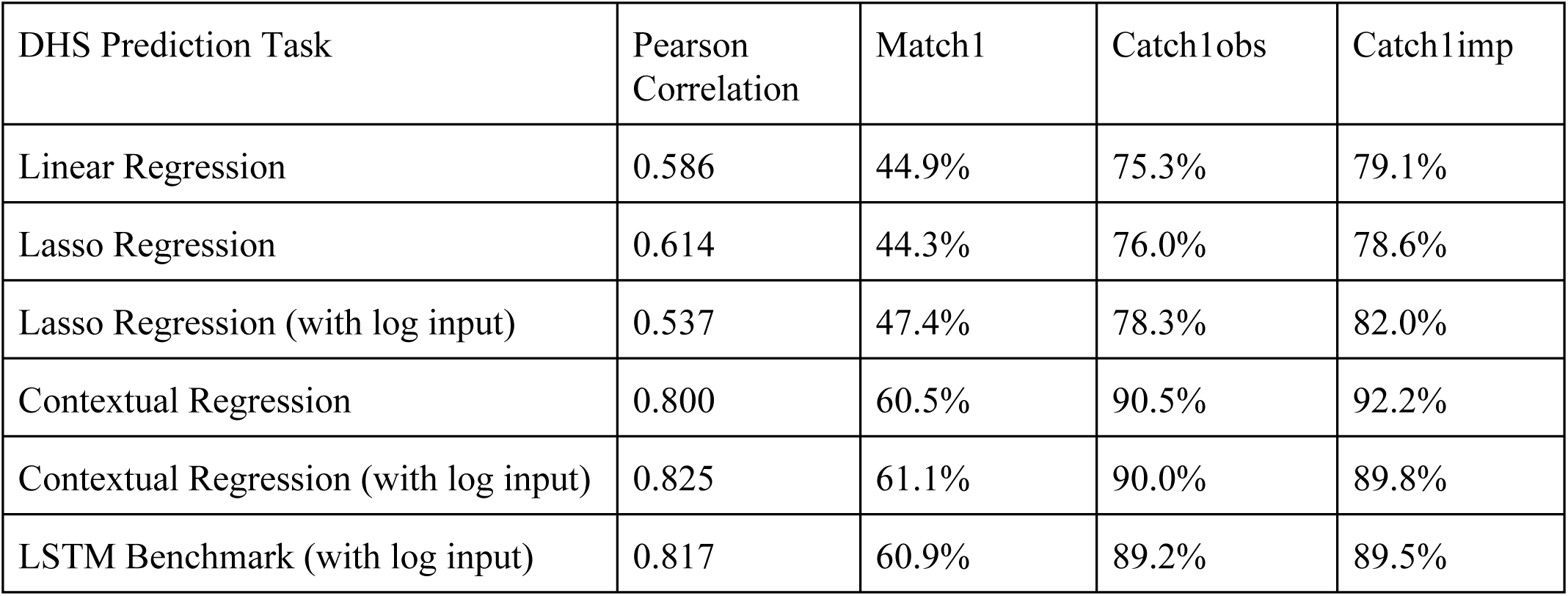
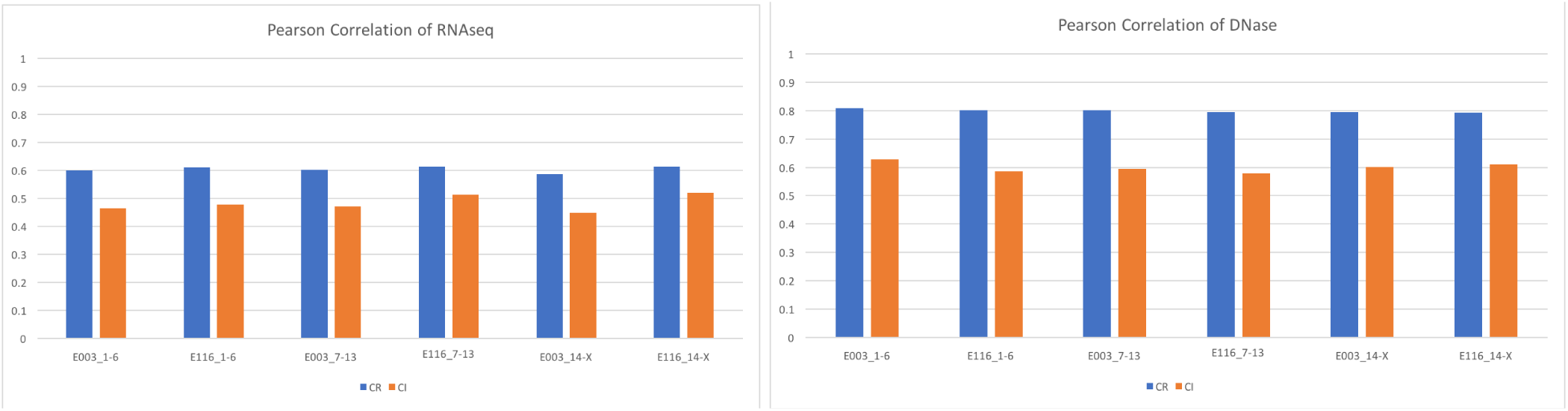
Prediction performance comparison of contextual regression with linear regression, LSTM model and ChromImpute-Top: Performance comparison of contextual regression with various linear regression and LSTM models on DHSs dataset in cell-line H1. We used four imputation quality evaluation metrics^28^: pearson correlation, match1 score (percent of 99 percentile experimental values in 99 percentile predicted values), Catch1obs (percent of 99 percentile experimental values in 95 percentile predicted values) and Catch1imp (percent of 99 percentile predicted values in 95 percentile experimental values). Bottom: Comparison of prediction accuracy (Pearson Correlation) between contextual regression (CR) and ChromImpute (CI).

Overall, the high prediction accuracy (relative error < 0.9% in all noise levels) and high rule extraction fidelity (RMSCD < 0.05 in all noise levels smaller than 1000%) on the simulated dataset support that our algorithm is highly reliable even under high noise level. These results also suggest that our method can be applied to datasets with missing information, as long as the effect of missing information on the prediction target is similar to a random noise of uniform distribution.

### Evaluating the Contextual Regression model on DNaseI hypersensitivity data

To examine the method in real experimental data, we applied it to predict DNaseI hypersensitivity sites (DHSs) using histone modifications in the H1 and GM12878 cell lines. We binned all the histone modification ChIP-seq and DHS-seq data at 200bp. To consider distal relationship, we included signals from 10kbp upstream to 10kbp downstream of the prediction location, i.e. 100 bins in total. In the H1 cells, there were 27 histone marks and thus the number of features reached 100 * 27 = 2700 which is too many for modeling. Hence, we arranged the features into a 100 by 27 matrix and factorized it into the tensor products of two vectors:

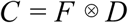

where C is the 100 * 27 combination feature matrix, F is the histone feature vector of length 27, corresponding to the number of histone marks, and D is the distal feature vector of length 100, corresponding to the number of bins. This approach greatly reduced the number of parameters in the model. In the current proof of concept stage, we further simplify and speed up the model by forcing the vector D to be the same for all the regions and applying a Lasso^27^ penalty on F.

We compared contextual regression with linear regressions and a benchmark neural network model implemented with Bidirectional-LSTM. The task for comparison is the prediction of DHS confidence values in cell line H1 using the 27 histone marks ChIP-seq data in the same cell line (Figure 3 and 4S). The models are trained on 70% of the randomly selected genomic regions and tested on the rest 30%. The contextual regression obviously outperformed linear regression, showing its better ability to capture nonlinear relationship between histone marks and open chromatin. At the same time, its performance is as good as the LSTM benchmark, showing that our interpretation method preserves the accuracy of a complex nonlinear model in this task.

**Figure 4.**
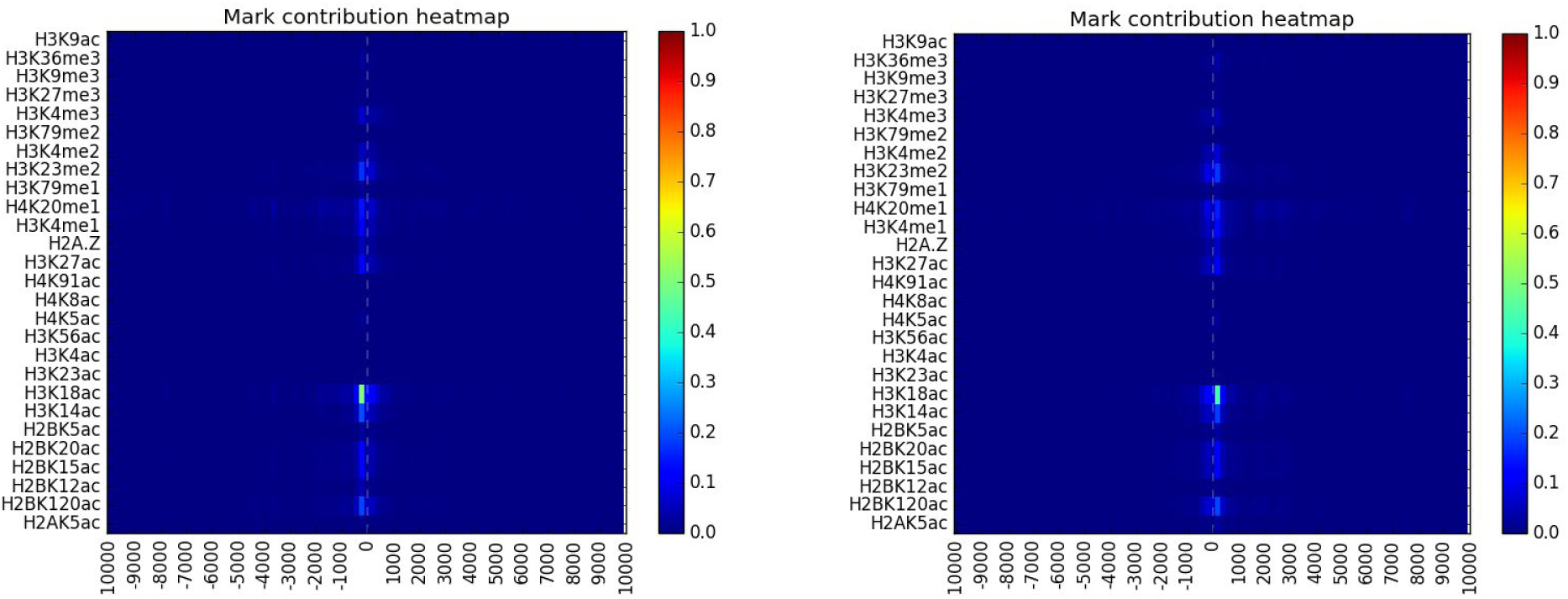
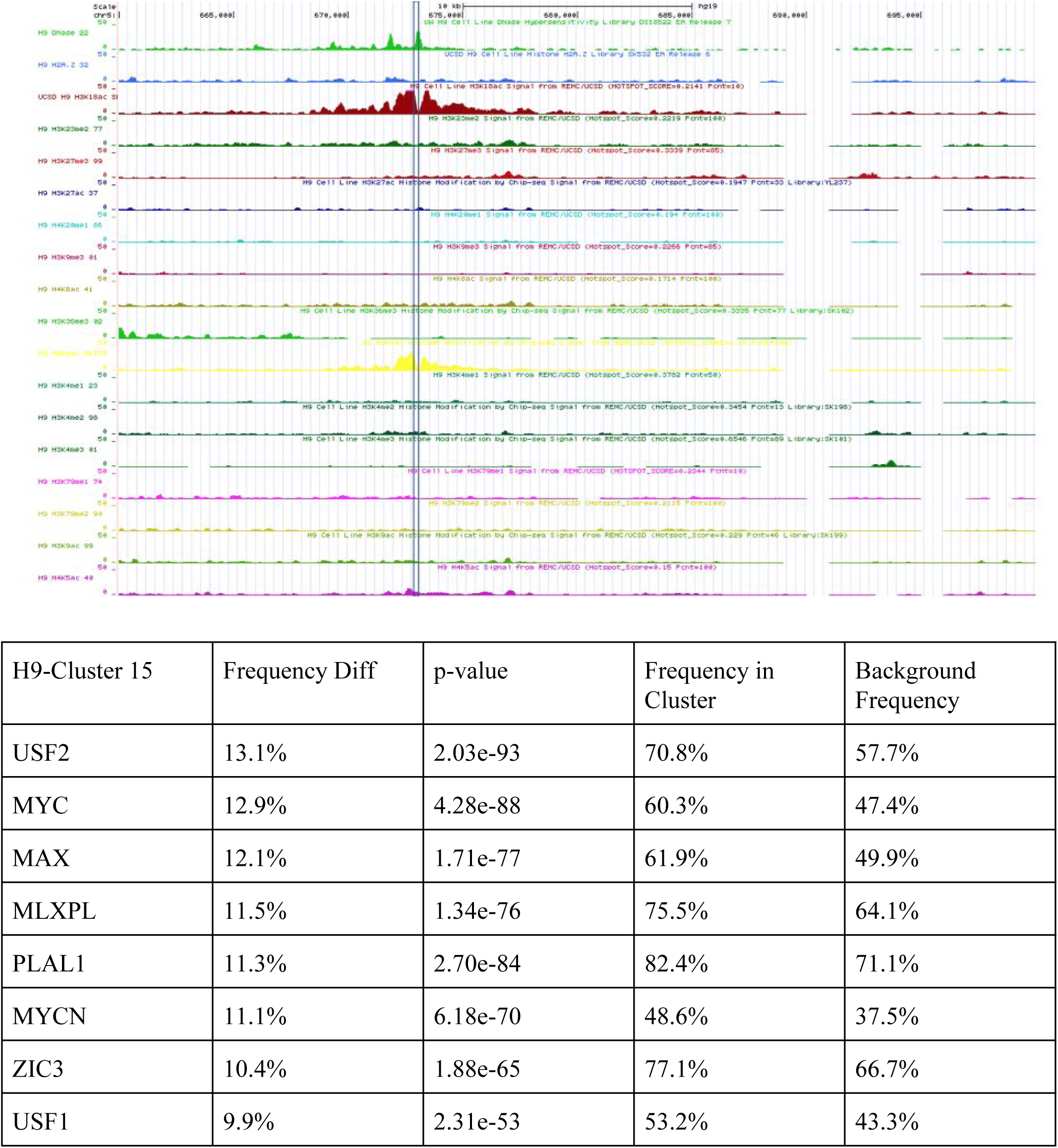
Study of H3K18ac pattern- Top: Mark contribution patterns of cluster 15 and 16 in H9 (H3K18ac pattern). Middle: Example genome browser view of the cluster 15 (H3K18ac dominant cluster). Bottom: Motif enrichment profile of cluster 15 in H9 (H3K18ac pattern).

Since applying log to the histone mark data improves the result slightly, we used this technique in all the studies in this section. We then assessed the performance of contextual regression on predicting RNA-seq and DHS data in cell lines H1 and GM12878 by comparing to ChromImpute^28^, the state of the art imputation method based on random forest algorithm. We ran both methods in three rounds of cross validations: (1) train: chr1-6, test: 7-X, (2) train: chr7-13, test: 1-6+14-X and (3) train: 14-X, test: 1:13. The models were trained using only the data from the same cell line. Contextual regression outperformed ChromImpute on both DHS and RNA-seq predictions (Figure 3 and 5S).

We also performed several other evaluations of contextual regression (See supplement for detail) on: high contribution feature validity, context weight assignment consistency and whether the target can be predicted well using only the high contribution features. As expected, the high contribution histone marks for both RNA-seq and DHS concord with previous research. Our model also assigned consistent weights to the features and made accurate prediction using only the high contribution features. These results have demonstrated that our method is not only able to achieve better performance than the state of the art models but robustly select important features as well.

### Discovering Important Histone Mark Patterns in the Open Chromatin Regions of H1

Next, we modified the model to search for both distal and histone mark features in the open chromatin regions in the H1 human embryonic stem cells. We made three changes to the model and training process: 1. we removed the restraint on distal feature vector D so that our model produces different D from different feature vectors input, 2. we did not apply log to make the result closer to the representation of the original magnitude, and 3. we applied a softmax^29^ constraint on D rather than the Lasso penalty on F to force the distal weights to be concentrated on a small number of locations. To make sure the patterns we extract are valid, we checked the prediction accuracy on the training and testing set and indeed, the model can still perform prediction accurately (Table 3S).

When analyzing the histone mark contribution patterns, we extracted the correctly predicted peaks (logpval > 3 for both experimental and predicted) from both training and testing set. We applied element-wise multiplication to the context weights (100 * 27) with the magnitude of the corresponding input (also 100 * 27), normalized them into vectors (of dimension 100 * 27 = 2700) of length 1, applied PCA (of dimension 2700), picked the top 100 PCs (which captured > 90% variance in all cell lines) and then clustered it with K-means clustering^30^. With k = 20, the clusters are well separated indicating by the great difference between in and across cluster distance from the cluster center (Figure 6S).

Among the cluster centers, we observed 4 types of major patterns: 1. the central dominant histone marks (with small contribution from other marks), 2. the spread histone marks, 3. the central dominant H2A.Z, and 4. the H3K27me3 pattern. Type 1 pattern (Figure 7S) has the peak contribution exactly from the center with a radius of 300bp by the H3K4me1/2/3 and H3K27ac. This type is composed of majority of the clusters (0, 1, 2, 3, 5, 6, 7, 8, 9, 12, 13, 17, 19). Type 2 patterns (Figure 8S) have long range contributions (> 300bp) besides the central dominant peaks, including clusters (11, 14, 15, 16). Type 3 pattern (Figure 9S) has central dominant (< 300 bp) contribution from H2A.Z, including cluster 10 and 18. Type 4 pattern (Figure 12S) has most significant contribution from H3K27me3 and ancillary contribution from H2A.Z, H4K8ac, H3K18ac and the H3K4me1/2/3. The effect of H3K27me3 is long range in this pattern and almost covers the whole 20kbp region. This type compose of cluster 4, a total of 3959 (4.2%) DNH peaks (logpval > 3 for both experimental and predicted).

Most of the open chromatin contributors found by our method, H3K27ac^31,32^, H2A.Z^33,34^, H3K4me1/2/3^35^, H4K20me1^36^ and K4K8ac^37^, have been repeatedly reported by previous research. However, H3K27me3 is not widely known to be associated with open chromatin, and their correlation with open chromatin is only mentioned in several papers^38,39^. To ensure this pattern is not caused by artifact, we visually inspected a couple of regions of type 4 pattern. All these regions have strong H3K27me3 signal that covers the whole region (Figure 10S). This is consistent with the contribution pattern (Figure 12S) discovered by our algorithm. We also found that the regions are enriched with transcription start sites (TSSs): about 64.4% of the regions in this cluster contain TSSs in 3kbp radius and 78.2% in 5kbp radius (Table 4S). Although visual inspection shows overall high concordance between open chromatin peaks with H3K4me2/3, there are regions that the DHS and H3K4me2/3 do not overlap well and H3K27me3 shows strong signals in the broad neighbor region. The most significant contribution from H3K27me3 to DHS peaks is surprising but consistent with the previous work^40^ which reports the bivalent domains that are stem cell specific: regions contain H3K4me3 and H3K27me3 sites of size 1kbp-18kbp and overlap with genes and TSSs. When applying our method to the DHS data in other cell-lines (H9, IMR90, GM12878, HUVEC) to search for cell line specific patterns, we found that pattern type 1-3 appear in all 5 cell-lines, however, type 4, the H3K27me3 pattern, is only found in H1 and H9, both are embryonic stem cells. This observation again coincides with the previous finding that the correlation between H3K27me3 and open chromatin is highly cell specific^38^ and mostly found in embryo cells^40,41^

### DNA motifs associated with open chromatin regions predictive by different histone marks

We next investigated whether there are specific DNA sequence motifs associated with each pattern. We compared the appearance frequency of motifs in each cluster and in all the clusters. The p-value in the comparison was calculated using One-Proportion Z-test. We limited our discussion only to cell-lines H1 and H9, which has the most diverse histone marks.

The cluster 4 and 16 of H1 and cluster 0, 4, 5, 8, 15, 16 and 19 of H9 contain motifs that have frequency difference above 10%. H9-8 and H9-19 are classic H3K27ac patterns. Both clusters have very similar motif enrichment in top 7 (Table 5S). Their enriched motifs agree except for NKX21 and PO3F1. Among the motifs enriched in both clusters, POU5F1 (Oct4) and SOX2 form a complex and regulate the expression of embryonic genes^42^. POU2F1/2 activate the genes that codes for H2B proteins^43^. POU3F2 is a protein involved in differentiation^44^. We also examined the H3K27ac dominant clusters in H1 (cluster 1 and 6) and found that many of the above motifs are also significantly enriched (Figure 11S and Table 6S). Thus a consistent motif enrichment pattern is confirmed for H3K27ac dominant open chromatin regions.

H1-cluster 4 and H9-cluster 4 are both H3K27me3 patterns (Figure 12S). Their motif enrichment profile are also very similar. Specifically, in H9-cluster 4, the frequency enrichment that is highly significant are shown in Table 1. In this cluster, MBD2 is the most enriched motif (62.9% overall -> 89.7% in cluster 4) and it plays the role of a transcription repressor that can recruit deacetylase and methyltransferase^44–48^. Another class of motifs that are enriched are the E2 family factors (E2F1-4) which are very important for the cell cycles^49,50^. These functions are highly correlated with the role of stem cell: these regions present in the differentiation process and can be an example of repressor induced chromatin remodeling^51^.

**Table 1.** Motif enrichment profile of cluster 4 in H9 (H3K27me3 pattern), pval of 0 values are caused by number underflow since the pval is smaller than machine accuracy

| H9-Cluster 4 | Frequency Diff | p-value | Frequency in Cluster | Background Frequency |
| --- | --- | --- | --- | --- |
| MBD2 | 26.8% | 0.0 | 89.7% | 62.9% |
| E2F1 | 25.6% | 0.0 | 83.2% | 57.6% |
| E2F4 | 24.3% | 0.0 | 63.5% | 39.2% |
| NRF1 | 23.8% | 0.0 | 75.9% | 52.0% |
| E2F3 | 21.5% | 0.0 | 37.1% | 15.6% |
| E2F2 | 15.0% | 1.75e-258 | 31.5% | 16.5% |

More interestingly, in H9 we found two marks, H3K18ac and H3K23me3, strongly associated with open chromatin that has not been reported. H9-cluster 15 and 16 are H3K18ac dominant clusters (Figure 4). The major contribution of H3K18ac is around 200bp up or downstream. The correlation of this mark with open chromatin is previously undiscovered and our algorithm only detected it in H9 among the 5 cell lines we studied. They compose 9.7% of the total predictable peak regions. To confirm the finding, we manually inspected some of the regions and H3K18ac does show up strongly around peaks of its prediction, although in some cases, the other marks also co-appear at the neighboring locations (Figure 4 and 13S). The motifs in the regions of cluster 15 and 16 show a unique enrichment profile (Figure 4). Among the enriched motifs, USF1,2 are associated with gene activation^52^. MYC family (MYC, MYCN, MAX and MLXPL) can form dimers and binds to E-box that regulates cell proliferation and differentiation^53^ while PLAL1 also facilitates the same function^54^. ZIC3 works on early body axis formation^55^. All these factors fit the stem cell nature of H9 cell line.

H9-cluster 5 is H3K23me2 dominant (Figure 5), which is another previously undiscovered and H9 unique pattern. The major contribution of H3K23me2 is around 200bp downstream. It compose 5.6% of the total predictable peak regions. However, the visual correlation of H3K23me2 with open chromatin is not as strong as the dominant marks in aforementioned clusters and co-appears with other marks in the open chromatin regions (Figure 5 and 14S). This lower correlation is also found statistically by our model, which shows less percentage of contribution from H3K23me2 (max value 43.8%) than the dominant marks in other clusters (for instance, max value of H3K18ac is 112%, > 100% since there are negative contributions from some other marks). The signature motif profile of H3K23me2 is very similar to the one of H3K27me3, with high MBD2 and E2 family enrichment. The largest motif frequency difference (Figure 5) is that the H3K23me2 regions has CEBPD motif frequency close to background level compared to the H3K27me3 cluster (81.2% vs 65.0%). CEBPB and CEBPG motif frequencies also show large difference between these two clusters. Function-wise, CEBPD is associated with immune response, immune cell activation and differentiation^56^. Other motifs that have significantly different profile among these two clusters is the E2 family. One possibility is that H3K23me2 is important for the stem cell development at a certain stage which is the reason why it is only enriched in certain open chromatin regions in H9.

**Figure 5.**
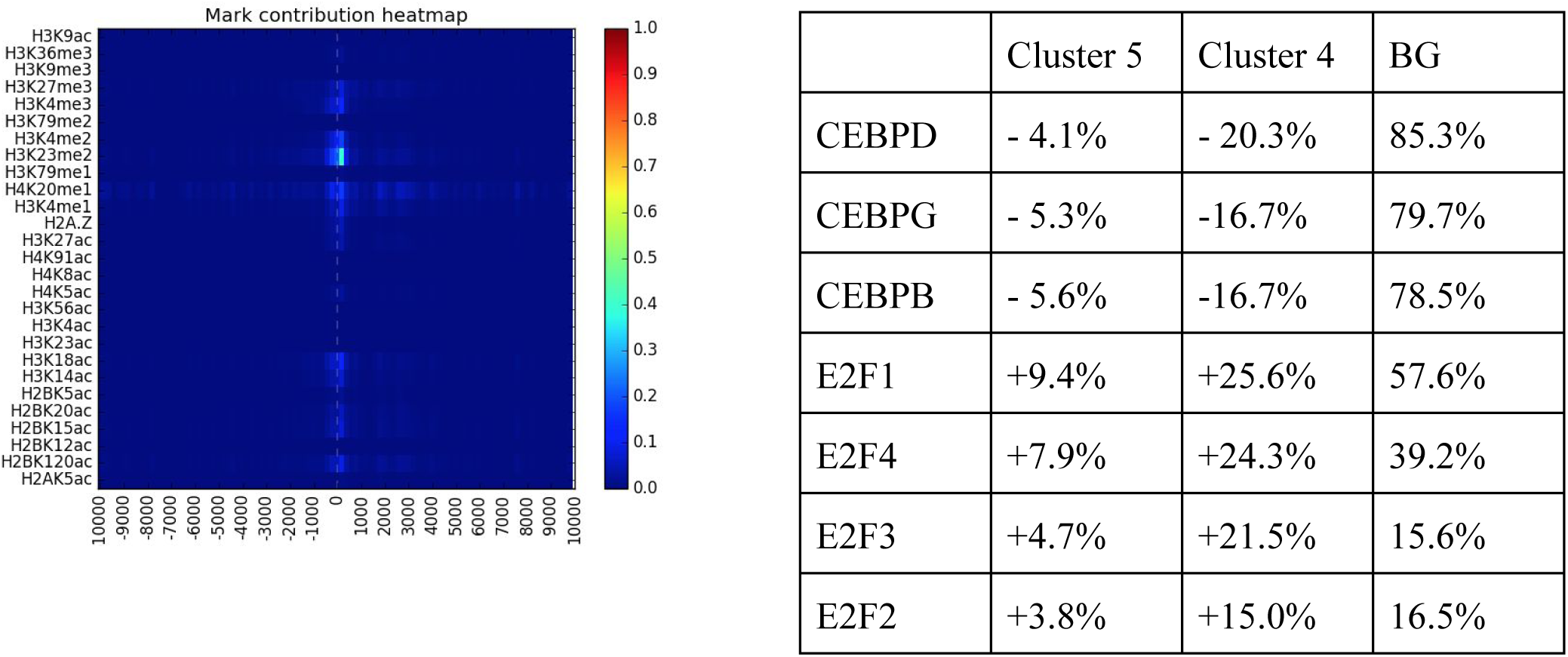
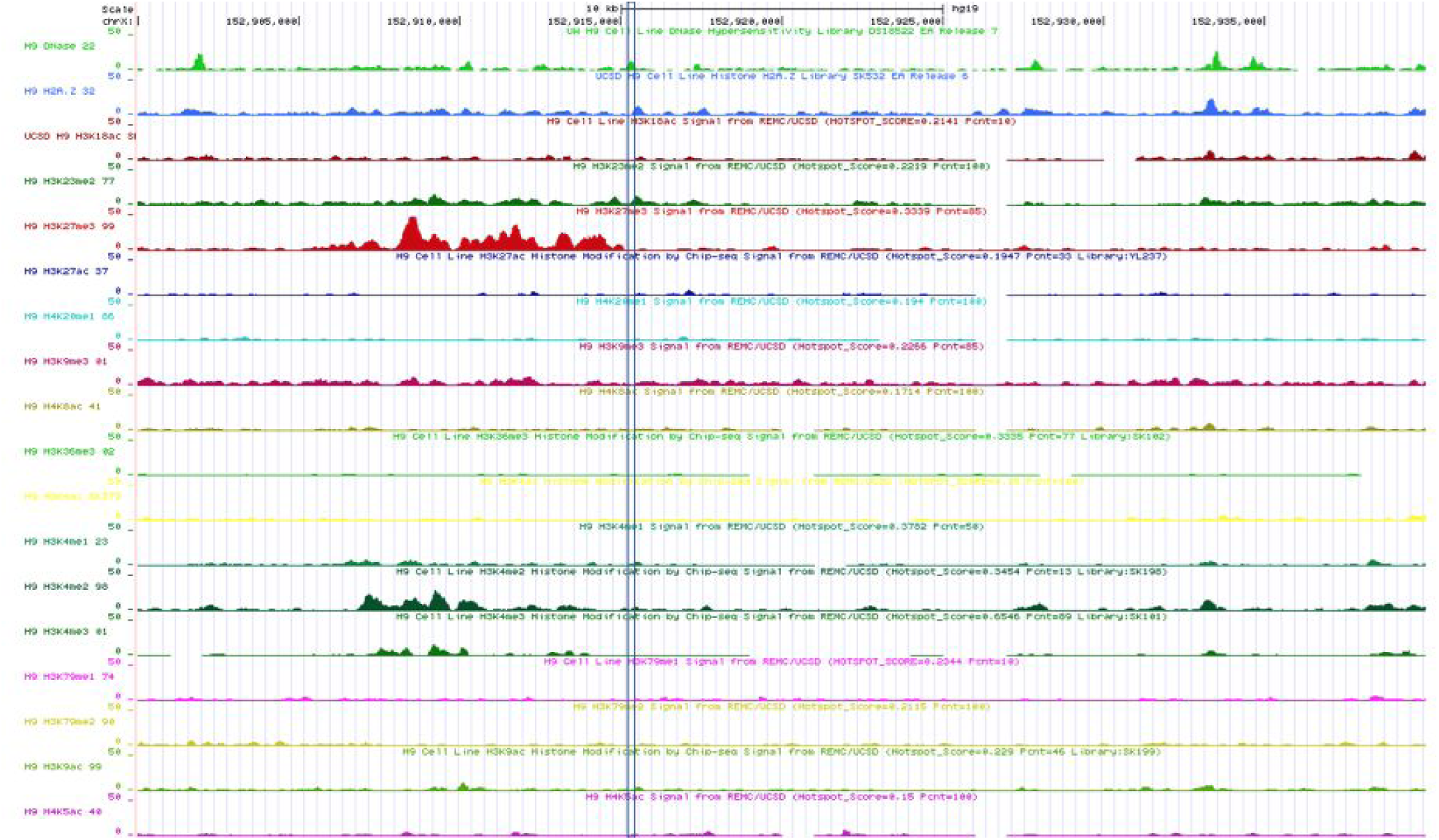
Study of H3K23me2 pattern- Top Left: Mark contribution pattern of cluster 5 H9 (H3K23me2 pattern), Top Right: Motif enrichment profile comparison between cluster 4 (H3K27me3 dominant) and 5 (H3K23me2 dominant) in H9. The 2nd and 3rd column are the motif frequency difference from the background and the 4th column is the BG (background) frequency, Bottom: Example Genome browser view of the cluster 5 (H3K23me2 pattern)

Furthermore, H1-cluster 16 and H9-cluster 0 (Figure 15S) are the spread type which include contribution from a lot of histone marks while none of them are dominant (Figure 16S). These two clusters show very similar motif enrichment patterns: strong contribution from MBD2 (+18%) and E2 (+5%-10%) families and high CEBPD frequency (more than 80% in both clusters). We hypothesize that this cluster is intermediate in its motif and histone mark patterns between the H3K27me3 and H3K23me2 clusters, which suggests that these three clusters are open chromatins in repressive states at different levels.

The above observations demonstrate that the open chromatin patterns captured by our method not only accord with visual inspection but enriched in unique motifs as well. This shows the existence of diverse histone mark and motif patterns that can relate to open chromatin via different mechanisms. Two of the patterns, H3K18ac dominant and H3K23me2 dominant have not been found in previous literatures, which shows the sensitivity of our method in the task of important feature detection.

## Discussion

In this paper, we described contextual regression, a generalized nonlinear model that is human interpretable. This method performs well in terms of prediction accuracy and important feature extraction in both simulated and experimental datasets. On epigenetic datasets, our method not only outperformed previous methods in terms of accuracy, but also robustly extracted important features that are aligned with previous research.

Using our method with the assistance of K-mean clustering, we not only found open chromatin patterns that have been discovered by previous research, but also new ones that exhibit signature patterns in both significant histone marks and DNA sequence motifs. H3K18ac, H3K23me2 and H3K27me3 are unfound or rarely mentioned indicators, which highlight the sensitivity of our method and the advantage of predictive models in the search of important factors. These results prove the validity of our algorithm and also emphasize the necessity of individual studies of each cell line, which will demand interpretable machine learning methods that can reduce the manual work needed in such kind of data mining tasks.

Despite its impressive performance, one potential problem of our method is the existence of alternative models in dataset with redundancy, which indeed exists in the open chromatin dataset. We have developed utopia penalty technique as a straightforward solution: adding a penalty term for feature weight unbalance during training. However, more research is needed to study the behavior of contextual regression to make it more stable and effective when dealing with this kind of situations.

